# Actomyosin dynamics, Bmp and Notch signaling pathways drive apical extrusion of proepicardial cells

**DOI:** 10.1101/332593

**Authors:** Laura Andrés-Delgado, Alexander Ernst, María Galardi-Castilla, David Bazaga, Marina Peralta, Juliane Münch, Juan Manuel González-Rosa, Federico Tessadori, Jeroen Bakkers, José Luis de la Pompa, Julien Vermot, Nadia Mercader

## Abstract

The epicardium, the outer mesothelial layer enclosing the myocardium, plays key roles in heart development and regeneration. During embryogenesis it arises from the proepicardium (PE), a cell cluster that appears in the dorsal pericardium close to the venous pole of the heart. Little is known about how the PE emerges from the pericardial mesothelium. Using the zebrafish model and a combination of genetic tools, pharmacological agents and quantitative *in vivo* imaging we reveal that a coordinated collective movement of the dorsal pericardium drives PE formation. We found that PE cells are apically extruded in response to actomyosin activity. Our results reveal that the coordinated action of Notch/Bmp pathways is critically needed for apical extrusion of PE cells. More generally, by comparison to cell extrusion for the elimination of unfit cells from epithelia, our results describe a unique mechanism where extruded cell viability is maintained.

---

The epicardium is the outer mesothelial layer of the heart. During development, the epicardium sustains the underlying myocardium through paracrine signals that promote its growth^1,2^. The epicardium is also an important cell source during embryogenesis. Epicardial-derived progenitor cells (EPDCs) differentiate into cardiac fibroblasts as well as other cell types including adipocytes^3^, mesenchymal stem cells, and smooth muscle or endothelial cells of the coronary vessels, and contribute to the formation of the annulus fibrosus and the valves^4,5^. After heart injury, EPDCs are involved in several aspects of tissue repair and regeneration, contributing to cardiac fibrosis, controlling the inflammatory response, promoting neoangiogenesis and cardiomyocyte proliferation^6^.

The epicardium derives from the proepicardium (PE), a cluster of cells that emerges from the pericardium close to the venous pole of the heart tube around the time of heart looping and after the onset of heart beating^7,8^. In the zebrafish, PE formation is regulated by bone morphogenetic protein (Bmp) signaling. Accordingly, mutants for the Bmp receptor acvr1l do not form a PE, whereas Bmp2b overexpression extends PE marker gene expression^9^. During the process of PE formation, cells undergo a change in polarity, suggesting that an epithelial-mesenchymal-like transition has a role in cluster generation^10–13^.

After the PE is formed, the heartbeat has an essential role in allowing PE cells to be “washed away” into the pericardial cavity. Heartbeat generates a pericardial fluid flow allowing the PE cells to reach the myocardial surface, to which they ultimately adhere and begin epicardial layer formation^14,15^. Nevertheless, while it is clear that pericardial flow triggers the release of PE cells to the pericardial cavity and is needed to form the epicardial layer, less is known about the role of mechanical forces on PE formation.

During morphogenesis, cell migration and proliferation results in the continuous rearrangement of mechanical properties of tissue layers. Collective cell migration and proliferation can lead to local cell crowding and the generation of tissue tension^16^. Conversely, changes in tissue growth can further influence cell signaling^17,18^. The actomyosin cytoskeleton plays a central role in controlling cell shape and developmental events^19–22^. It is tightly associated with membrane junction complexes and can react to extracellular signals or signals from neighboring cells by altering cell properties^22–24^. From looping, trabeculation to valvulogenesis, mechanical activity of the heart has a fundamental function in controlling cardiac morphogenesis^25–28^. Here we used the zebrafish model to study the morphogenetic events leading to PE formation. We found that cells from the dorsal pericardium (DP) collectively move towards the DP midline, where some of them round up and extrude into the pericardial cavity. This movement of DP cells to the midline and PE cell extrusion is dependent on actomyosin dynamics. We found that Bmp2 signaling lies upstream of the actomyosin cytoskeletal rearrangements necessary for PE formation. Furthermore, we show that the developing heart tube influences PE formation as endocardial Notch signaling controls Bmp expression.

## RESULTS

### Constriction of the dorsal pericardium and apical extrusion leads to PE delamination

To investigate the formation of the PE, we performed a detailed analysis of pericardial mesothelial cell movement in zebrafish embryos. PE clusters appeared at the midline of the DP, which extends from the venous to the arterial pole of the heart tube (Fig. 1a). To image PE formation, which begins around 52 hours post fertilization (hpf), we used the enhancer trap line *Et(−26.5Hsa.WT1-gata2:EGFP)^cn1^* (hereafter termed epi:GFP) in which GFP expression is controlled by the regulatory elements of *wilms tumor 1a* (*wt1a*), and recapitulates its expression pattern^14^. Thus, pericardial and PE cells are GFP^+^ in the epi:GFP line. GFP expression is present in all DP cells and is particularly strong around the cell nucleus^29^ and thus suitable for cell tracking. We tracked individual cells of the DP using time-lapse video imaging from 52 to 60 hpf, and found that they became displaced and converged at the midline (Fig. 1b,c and Supplementary Movie 1). We also measured the angle of the cell trajectories of the constricting DP tissue in relation to the midline and found that the majority were close to 90°, indicating that DP cells move nearly perpendicular to the midline and suggesting that this movement leads to an active directional accumulation of cells at the midline producing DP tissue constriction (Fig. 1d; n=3 embryos).

**Fig. 1.**
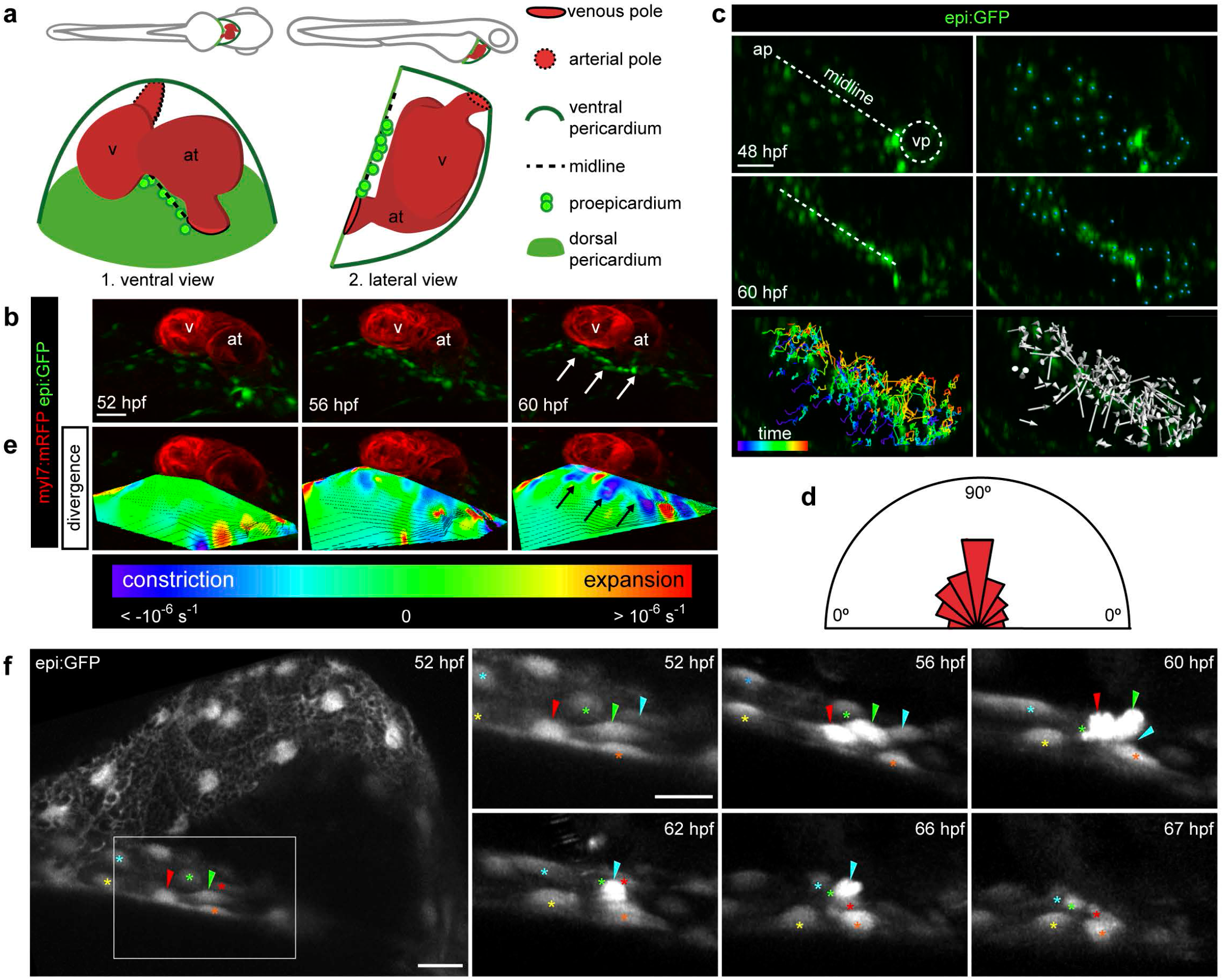
Collective cell movements and apical extrusion lead to PE delamination. **(a)** Scheme of lateral view of a 60 hpf zebrafish embryo. Zoom on heart below overview. Lateral view is rotated by −90°. Legend for structures are on right side of the panel. Shown are the dorsal and ventral pericardium, ventricle, atrium, outflow tract venous pole, midline and proepicardium (PE). **(b)** 3D ventral view of the heart region in a *epi:GFP; myl7:mRFP* double transgenic larvae. epi:GFP^+^ cells in the dorsal pericardium (DP) and PE are shown in green, *myl7:mRFP* in red (myocardium). Maximal projection from an *in vivo* time-lapse at different time-points is shown. White arrows point to PE clusters (see also Movie 1). **(c)** First and last frame of an *in vivo* time-lapse DP cells from an *epi:GFP* larvae; the midline is shown by a discontinuous white line. Blue dots indicate the tracked cells. Full colored tracks label the first time frame in purple and the last in red. Arrows indicate overall direction of tracked cells towards the midline. **(d)** The half-rose diagram shows the number of DP cells from 3 animals for which the angle of the track was measured relative to the midline. **(e)** 3D reconstruction of the divergence field of the DP on top of images from b. Blue regions represent constriction and red expansion of the tissue. Black arrows point to PE clusters (see also Movie 2). **(f)** epi:GFP *in vivo* time-lapse. A maximum intensity projection of 22 μm is shown. Left panel shows an overview of the vp and DP at 52 hpf. Right panels show zoomed frames of the time-lapse from 52 to 67 hpf. Colored arrowheads indicate emerging PE cells. DP cells surrounding emerging PE cells are labeled with colored asterisks (shown is 1 out of 10 acquisitions, see also Movie 3). ap, arterial pole; at, atrium; hpf; hours post fertilization; v, ventricle; vp, venous pole. Scale bar: 50 μm (in **f** overview 20μm and time-lapse 10μm).

To further characterize the morphological changes to the DP during PE formation, we quantified the changes in the relative velocity of DP cells with respect to their neighboring cells using a customized calculator of the velocity field divergence based on DP cell tracking in epi:GFP embryos (Supplementary Fig. 1). The divergence field represented predominantly by purple-blue colors indicates an overall constriction in the tissue, whereas tissue expansion is reflected by red-orange pseudostaining within the divergence field. By calculating this factor for the imaged DP at 43 different time points every 12 min, and measuring 150 DP cell tracks in three embryos, we confirmed that there was an overall constriction of the DP from 52 to 60 hpf, with highest local levels of constriction at the site of PE formation (Fig. 1e and Supplementary Movie 2).

We next characterized the emergence of PE cells, again by *in vivo* imaging of the epi:GFP line. During the displacement of DP cells towards the midline, cells close to the midline began to show a stronger GFP signal and rounded up (Fig. 1f and Supplementary Movie 3, which is representative for the event observed in a total of 10 different movies). We further observed that the cells surrounding these emerging PE cells came closer together. Ultimately, one or multiple cells protruded from the DP layer and remained only loosely attached to the neighboring DP mesothelial cells. The cells that were bordering the newly formed PE cells subsequently converged under the rounded PE cell. These cell behaviors are landmarks of apical extrusion, allowing cells to bulge out and leave organized epithelia^30^. Therefore, the results suggest that PE formation occurs through the local overcrowding of cells at the DP midline, inducing apical extrusion of mesothelial cells of the DP.

### Proliferation of pericardial cells contributes to constriction at the midline

To determine whether the overall constriction of the DP was in part due to cell proliferation, we analyzed the spatial distribution of cell division within the DP. By measuring the distance of the cell division plane to a defined midline (a surface of about 20 μm diameter drawn from the arterial to the venous pole of the heart tube) we found that pericardial cells divided all over the DP (Fig. 2a,b and Supplementary Movie 4). However, cell division occurred neither directly at the midline nor preferentially close to it (inside the surface = 0 μm 0 divisions, 0–20 μm 7 divisions, 20–80 μm 11 divisions, Fig. 2c). Categorizing orientation of cell divisions showed that most divisions were perpendicular to the midline (Fig. 2d). We next assessed cell proliferation by immunostaining for phospho-histone 3 (pH3) on fixed epi:GFP embryos. While proliferating cells could be found in the pericardium (Fig. 2e-g and Supplementary Fig. 2a,b), only one pH3^+^ PE cell was observed in 4 out of 12 embryos (Fig. 2g,h Supplementary Fig. 2c,d). Inhibiting cell proliferation from 48 hpf onwards with the pharmacological agents nocodazole (noc) or aphidilcolin/hydroxyurea (aph/hydrU) significantly reduced the number of pH3^+^ cells in the pericardium from 52 ± 29 in controls to 21 ± 8 in noc and 0 ± 1 in aph/hydrU-treated embryos at 60 hpf; (Fig. 2e,f). Noc inhibits cell proliferation by halting mitosis^31^ and aph/hydrU inhibits DNA synthesis^32^. Both treatments significantly reduced the number of PE cells from 9 ± 3 to 0 ± 1 or 2 ± 1, respectively (Fig. 2i).

**Fig. 2.**
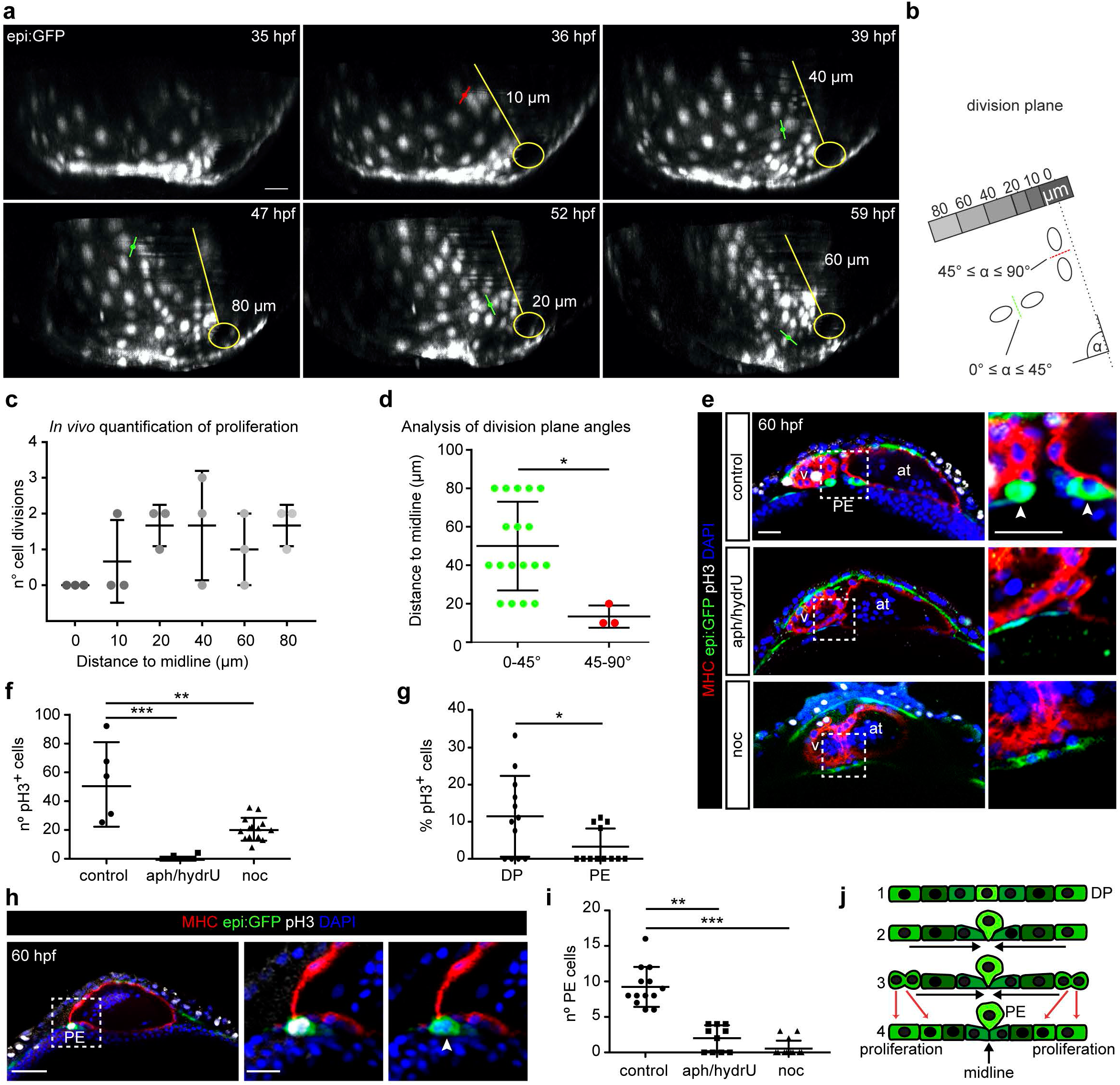
Proliferation of pericardial cells is necessary for PE formation. **(a)** Six frames of an epi:GFP *in vivo* time-lapse (see also Movie 4). The dorsal pericardium (DP) was isolated and presented from a top view. Division planes are indicated with red (45° ≤ α ≤ 90°) and green lines (0° ≤ α ≤ 45°). A representation of the midline is shown in yellow. Micrometer values show the measured distance of the division to the midline **(b)** Scheme showing the principle of measurements for distance and angles of division planes to midline. Graphs show number of divisions during *in vivo* time-lapse relative to the midline (3 embryos with 20 cell divisions; P-value= 0.2, unpaired Student’s T-test) **(c)** and angle of midline to division plane relative to distance to the midline **(d)** The division plane was more often perpendicular than parallel to the midline P-value= 0.0144, unpaired Student’s T-test. **(e)** Whole mount immunofluorescence with anti-GFP (green), Myosin Heavy Chain (red) and anti-pH3 (white). DAPI counterstains nuclei (blue). Shown are optical transversal sections of a control heart and hearts form larvae treated with the proliferation inhibitors aphidilcolin/hydroxyurea (aph/hydrU) or nocodazol (noc). Zoomed view of the dotted PE area is shown on the right. Arrowheads point to the PE. **(f)** Quantification of total number of pH3^+^ cells in the pericardium. **(g)** Percentage of pH3^+^ cells in the DP compared with the PE. **(h)** Zoomed view of a unique pH3^+^ PE cell in a control embryo. **(i)** PE cell number from conditions shown in e. **(j)** Scheme of apical PE extrusion mechanism: 1) DP is a flattened mesothelium; 2) DP cells move towards the midline; 3) PE cells rounds up at the midline; 4) proliferation of DP cells at the border site also contributes to the constriction and PE cells finally extrude. at, atrium; DP, dorsal pericardium; hpf, hours post fertilization; PE, proepicardium; v, ventricle. Scale bar: 50 μm (in **h** zoomed images, 20μm). All data are means ± s.d., one-way ANOVA followed by Kruskal-Wallis significant difference test was used in **f** and **g**; (Unpaired two-tailed Student’s *t*-test was used in panel i). * *P* < 0.05; ** *P* < 0.01, *** *P* < 0.001.

However, we observed that in noc-treated embryos, DP cells still converged at the midline (Supplementary Movie 5). Thus, cell proliferation might contribute partially to the local crowding of DP cells at the midline, which ultimately leads to PE cluster formation (Fig. 2j).

### PE formation depends on actomyosin dynamics

Cytoskeleton reorganization is a fundamental process for the apical extrusion of apoptotic epithelial cells and for cell migration/invasion of metastatic cells^33–35^. Prior to extrusion, cells present an overall increase in F-actin levels and a change in actomyosin localization, from basal to apico-cortical deposition^36^. Because myosin II plays a fundamental role in actin filament motion^37–39^, we analyzed its expression in PE cells. Immunofluorescence analysis revealed that myosin II-A was highly expressed in PE cells (Fig. 3a’). In DP cells close to the midline, myosin II-A expression was strong at the cell boundaries (Fig. 3a’’). In rounded PE cells, myosin II-A was also expressed apically and at boundaries between PE cell pairs (Fig. 3a’’’, 3a’’’’). *In vivo* imaging of the *actb2:myl12.1-mCherry^e1954^* line, expressing myosin II fused to mCherry under the embryonic ubiquitous promoter *actin beta 2* (*actb2*), confirmed an accumulation of myosin II in DP cells at the midline and in the PE cluster (Supplementary Movies 6,7).

**Fig. 3.**
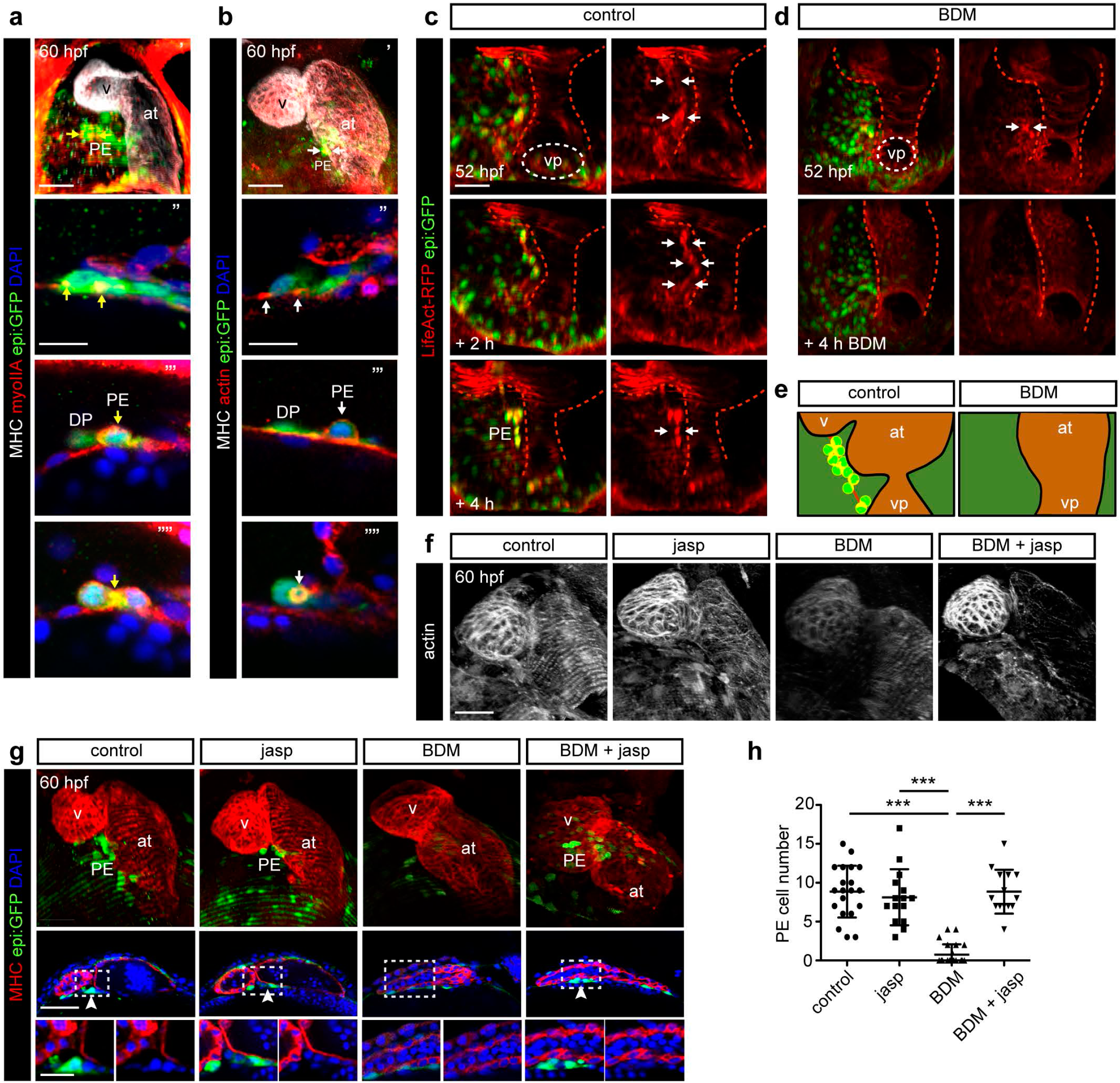
PE formation depends on actomyosin dynamics. **(a)** Upper panel shows a maximum projection of a 60 hpf zebrafish heart immunostained for Myosin Heavy Chain (MHC, white), myosin II-A (red) and GFP (green). Nuclei are counterstained with DAPI (blue). The three bottom zoomed images are different views of the PE region. Yellow arrows point to myosin II-A concentration sites. DP indicates a flat dorsal pericardial cell. **(b)** Immunostaining as in **a**, but with phalloidin-488 to visualize F-actin (red). White arrows point to actin concentration sites. **(c, d)** Ventral view of the heart region in a *epi:GFP; lifeActin-RFP* larvae at different time points of an *in vivo* time-lapse. Ventral pericardium and heart tube was post-process eliminated and the heart tube shape was drawn with a red discontinuous line. **(c)** Untreated control; white arrows mark sites of actin concentration in the DP. **(d)** BDM-treated larvae. **(e)** Scheme at 58 hpf of the actin concentration at DP and PE clusters during PE formation in untreated or BDM-treated larvae. Actin is in red, DP and PE cells in green. **(f)** Maximal projection of hearts labeled for actin in fish untreated or treated with jasp and BDM. **(g)** Top panels show a maximum projection of a 60 hpf zebrafish heart, and bottom panels show an optical section of epi:GFP animals immunostained for MHC (red) and nuclei counterstained with DAPI (blue). Larvae were either untreated or treated with combinations of jasp and BDM. White arrowheads indicate the PE. **(h)** Quantification of PE cell number in **g**. at, atrium; BDM, Butanedione Monoxime; DP, dorsal pericardium; hpf, hours post fertilization; jasp, jasplakinolide; PE, proepicardium; v, ventricle; vp; venous pole. Scale bar: 50 μm (in **a**“-**a**““4 and **b**“-**b**““is 10μm). Data are mean ± s.d., one-way ANOVA followed by Kruskal-Wallis significant difference test. \*\*\**P* < 0.001.

We also analyzed the localization of polymerized actin during PE cell formation at 60 hpf using whole-mount labeling of actin with a phalloidin-coupled fluorophore (Fig. 3b’). Whereas actin was located in the basal region of the DP cells (Fig. 3b’’, 3b’’’), it was polarized and accumulated apico-cortically in PE cells, consistent with the pattern observed for myosin II (Fig. 3b’’’,3b’’’’). In vivo characterization of actomyosin dynamics was performed in the double transgenic line *βactin:LifeAct-RFP^e2212Tg^*;epi:GFP. We observed that from 52 to 60 hpf, F-actin concentrated in DP cells at the midline where the PE appears (Fig. 3c). Indeed, a thick actin cable was visible in epi:GFP^+^ cells at the time of PE formation, spanning the midline of the DP (Supplementary Movies 8–11). Actin was concentrated basally in the DP cells that surrounded the PE cells, whereas in PE cells actin changed its polarity and and became localized cortically and concentrated in the contact region between the rounded up PE cells (Supplementary Movie 12). In PE cells that were close to being released into the cavity, we observed that in a final stage prior to release a thin actin rich stalk appeared in the PE cell still attached to the DP (Supplementary Movie 13). In sum, our data suggest that PE formation occurs concomitantly with extensive actin reorganization: before PE formation, F-actin is located basally in DP cells, then it becomes concentrated in PE precursor cells and it finally accumulates all around the cortex of the rounded PE cells. We previously found that PE formation is impaired in the presence of butanedione monoxime (BDM)^14^, which interferes with myosin-ADP-Pi phosphate release, and locks myosin II into a low affinity conformation with actin impeding myosin movement on top of actin filaments^40^. Inhibition of myosin II with blebbistatin (BLEB), through maintenance of myosin II in an actin-detached state, also impaired PE formation and this was dose dependent (Supplementary Fig. 3a-c). Compared to non-treated embryos, BDM treatment led to a reduction in LifeAct-RFP expression in the DP (Fig. 3d,e), showing that F-actin is unstable in DP cells at the midline. Results from *in vivo* imaging revealed that the arrangement of epi:GFP^+^ cells at the midline was inhibited by BDM and the movement of the DP was impeded (Supplementary Movies 14,15).

We reasoned that if actin polymerization was required for PE formation, pharmacological enhancement of actin stability should counteract the effect of myosin II inhibition. To test this, we administered jasplakinolide (jasp), which promotes actin filament polymerization and stabilization^41^, to epi:GFP animals in the presence or absence of BDM. Embryos treated with jasp and BDM from 48 hpf onwards showed stronger actin signals than embryos treated only with BDM, suggesting that jasp rescued actin polymerization in BDM-treated animals (Fig. 3f). Moreover, animals treated with jasp or BDM + jasp had 8 ± 4 and 9 ± 3 PE cells, respectively (n=15 or n=14 embryos each), which resembles control conditions, whereas BDM-treated animals had 1 ± 1 PE cells (n=13) (Fig. 3g,h). Thus, enhancement of actin filament polymerization and stabilization in the presence of BDM correlated with PE formation, suggesting that F-actin is necessary for PE formation.

### Bmp2b influences actomyosin dynamics during PE formation

During heart tube development, myocardial cells are involved in the secretion of signaling molecules such as Bmps^42–44^, and Bmp signaling is necessary for correct PE cluster formation in zebrafish^9^. To evaluate whether ectopic Bmp could rescue the impairment in PE formation upon inhibition of actomyosin dynamics, we next performed experiments using the transgenic line *hsp70:bmp2b*^45^ crossed into epi:GFP, which allows *bmp2b* levels to be increased at a specific developmental stage by heat shocking the embryos at 39°C for one hour. In control (non-transgenic for *hsp70:bmp2b*) animals subjected to heat shock (HS), the PE was completely formed and visible at 60 hpf and comprised approximately 10 cells (Fig. 4a,b). Overexpression of *bmp2b* during the embryonic stages preceding PE formation by delivering HS pulses at 26, 32 and 48 hpf, resulted in a 2-fold increase in the number of PE cells; from 10 ± 5 cells in control embryos to 22 ± 9 in bmp2b-overexpressing embryos at 60 hpf (P < 0.0001) (Fig. 4a,b). The total number of DP cells prior to PE formation did not differ between bmp2b-overexpressing fish and control animals heat shocked for 26 and 32 hpf (Supplementary Fig. 3d; n=6/7 animals), suggesting that the enlarged PE clusters observed upon *bmp2b* overexpression were not a consequence of an expanded population of DP cell precursors. PE cells emerge to a small extent from the DP close to the venous pole of the heart (vpPE) and to a larger extent from the DP close to the atrioventricular canal (avcPE)^14^. To assess whether Bmp2b was acting on a particular subpopulation we individually quantified the number of cells in the avcPE and vpPE clusters and found that *bmp2b* overexpression significantly increased the number of cells in the avcPE cluster (Supplementary Fig. 3e).

**Fig. 4.**
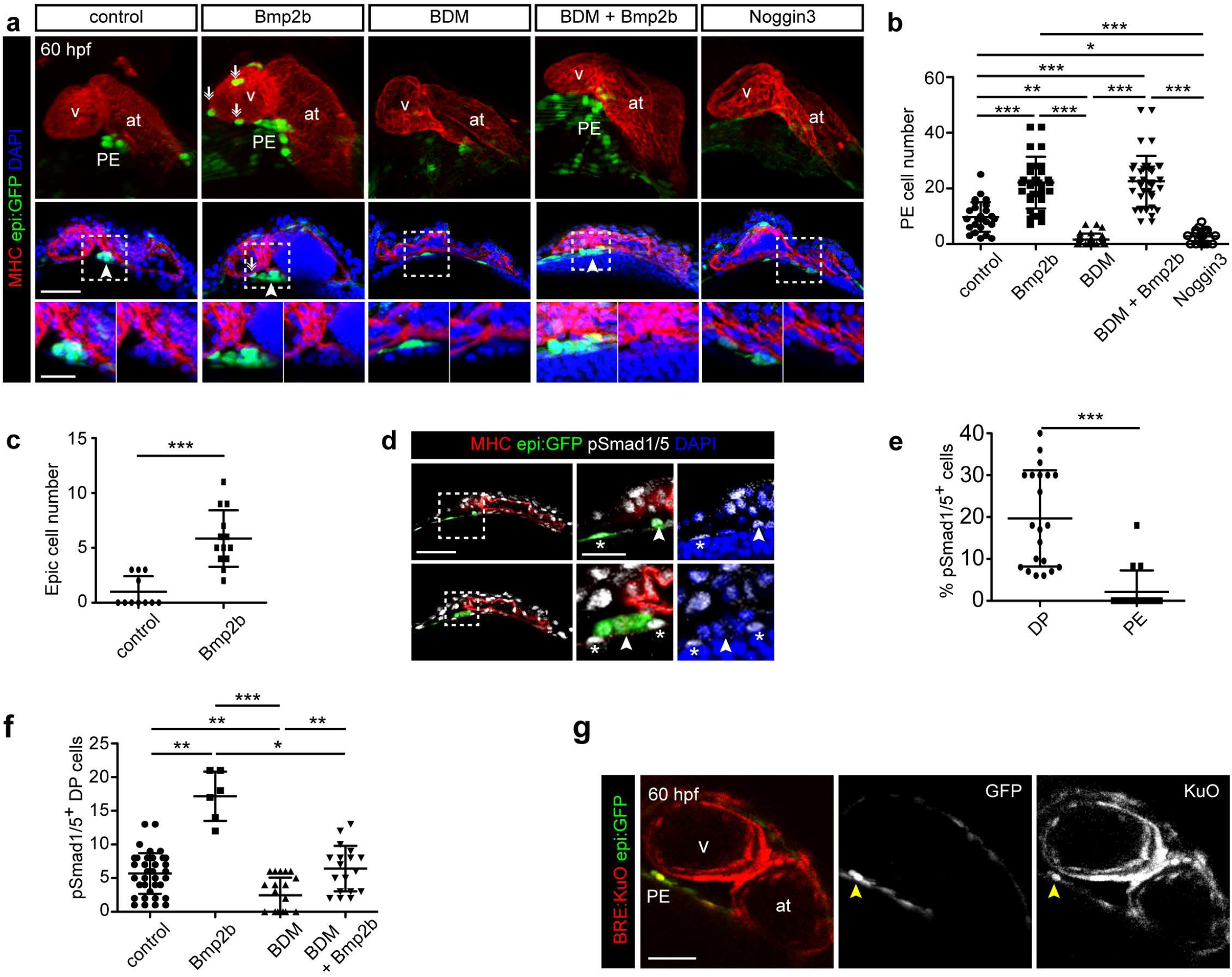
Bmp2b rescues PE formation upon Myosin II inhibition. **(a)** Top panels show a maximum projection of a 60 hpf zebrafish heart, and bottom panels show an optical section of epi:GFP animals immunostained for GFP in green and Myosin Heavy Chain (MHC) in red and nuclei counterstained with DAPI (blue). Untreated fish were compared with those overexpressing *bmp2b* with and without BDM or with those overexpressing *Noggin 3*. Arrowheads point the PE. Arrows mark epicardial cells. **(b)** Quantification of PE cell number in **a. (c)** Quantification of epicardial cell number at 60 hpf in bmp2b-overexpressed versus non-overexpressed animals. **(d)** Optical sections through epi:GFP hearts at 60 hpf showing pSmpad1/5 staining. Zoomed views are also shown. epi:GFP animals were immunostained for GFP in green, MHC in red, pSmad1/5 in white. Nuclei were counterstained with DAPI (blue). Arrowheads point to PE cells and asterisks mark DP cells. **(e)** Percentage of pSmad1/5+ cells in the DP compared to the PE. **(f)** Total number of pSmad1/5+ cells in the pericardium at 60 hpf. **(g)** Optical section from an *in vivo* time-lapse of an BMP reporter line *BRE:KuO/* epi:GFP embryo. Yellow arrowheads point to the double positive cells in the PE. at, atrium; BDM, Butanedione monoxime; DP, dorsal pericardium; epic, epicardium; hpf, hours post fertilization; PE, proepicardium; v, ventricle. Scale bar: 50 μm (in **a** and **d** zoomed images, 20μm). Data are mean ± s.d., one-way ANOVA followed by Kruskal-Wallis significant difference test. Unpaired two-tailed Student’s *t*-test was used in panel **c** and **f.** * *P* < 0.05; ** *P* < 0.01, *** *P* < 0.001.

In addition, the number of epicardial cells found on the ventricular myocardial surface was higher in bmp2b-overexpressing fish at 60 hpf (Fig. 4a,c). In line with this, antagonizing Bmp signaling by overexpressing *noggin3* in the *hsp70l:noggin3*^*fr1*^*4* line ^45^ through HS at 48 hpf significantly reduced the number of GFP cells within the PE cluster (Fig. 4a,b).

We next aimed to assess whether Bmp2b activated Bmp signaling directly in PE cells. To do this, we evaluated the expression of its downstream effector pSmad1/5. pSmad1/5 immunostaining was found both in DP cells and in some PE cells between 55 and 60 hpf (Fig. 4d-f). Approximately 20% of DP cells were pSmad1/5+ (a mean of 4 ± 3 positive cells from 15 ± 7 total DP cells; n=21 embryos), whereas only about 2% of PE cells were pSmad1/5+ (0 ± 1 positive cells from a total of 9 ± 4 cells; n=16 embryos). The number of pSmad1/5+ DP cells was significantly higher than the number of pSmad1/5+ PE cells (P < 0.001). Moreover, the total number of pSmad1/5+ DP cells was increased in bmp2b-overexpressing embryos (Fig. 4f). We futher used a Bmp reporter line expressing Kusabira Orange (KuO) under the control of a promoter harboring several smad binding sites, named Bmp responsive elements (BRE)^46^. Results from *in vivo* imaging of BRE:KuO; *epi*:GFP fish revealed that DP cells and some PE cells were KuO+ (n=1-2 KuO+ PE cells per animal, seen in 3 animals from 52 to 58 hpf), confirming that the Bmp signaling pathway was transiently active in PE cells (Fig. 4g and Supplementary Movie 16). Therefore, an increase in Bmp activity within the DP correlates with PE formation.

We addressed whether overexpression of *bmp2b* could rescue the impaired PE cluster formation with BDM from 48 hpf onwards. PE formation was observed in bmp2b-overexpressing animals treated with BDM, and clusters were larger than those observed in controls (23 ± 3 vs 10 ± 5 cells; Fig. 4a,b *P* < 0.0001; n=29 embryos). Overexpression of *bmp2b* also rescued PE formation in BLEB-treated animals (Supplementary Fig. 3a,b). We observed that the rescue of PE formation was dependent on *bmp2b* expression levels. Accordingly, larger PE clusters were detected after 3 HS pulses (at 26, 32 and 48 hpf) (22 ± 9 cells; n= 25 embryos) (Supplementary Fig. 3b) than with only one HS pulse at 48 hpf (10 ± 4 cells; n=30 embryos) (Supplementary Fig. 3d). We also assessed at which developmental stage the effect of bmp2b was more prominent on PE formation. A unique HS at 48 hpf rescued PE formation at 60 hpf in BDM-treated hearts (8 ± 4 cells vs 3 ± 2, n=20 embryos) (Supplementary Fig. 3f), but a single HS at 26 hpf failed to rescue PE formation at 60 hpf in BDM-treated embryos (6 ± 3 cells vs 4 ± 2, n=12 embryos) (Supplementary Fig. 3g). *bmp2b* overexpression after BDM treatment also failed to rescue PE formation (Supplementary Fig. 3h, n=10).

BDM treatment impairs pericardial cell proliferation^14^. As we observed that DP proliferation events preceded PE formation, we questioned whether *bmp2b* overexpression in BDM-treated embryos might increase DP cell proliferation. However, the number of proliferating epi:GFP^+^ pericardial cells was very low and not different between BDM-treated animals overexpressing or not *bmp2b* (Supplementary Fig. 2b,d, n=23). We therefore explored how the actomyosin network is altered by *bmp2b* overexpression to understand the mechanisms through which Bmp2b might neutralize the effect of BDM on the actomyosin cytoskeleton. BDM treatment strongly reduced the amount of myosin II-A in DP cells (n=8 animals) and overexpression of *bmp2b* in BDM-treated animals rescued the apical polarization of myosin II-A in PE cells (n=11) to levels similar to those observed in control embryos (n=10) (Fig. 5a), suggesting a recovery of actomyosin dynamics by the Bmp pathway. Thus, we investigated how actin polymerization is affected upon Bmp2b overexpression. Examination of F-actin levels by immunostaining at 60 hpf revealed that *bmp2b* overexpression increased these levels significantly. In the presence of BDM, actin levels in PE cells were lower than those in controls; however, they were significantly rescued upon *bmp2b* overexpression (Fig. 5b,c).

**Fig. 5.**
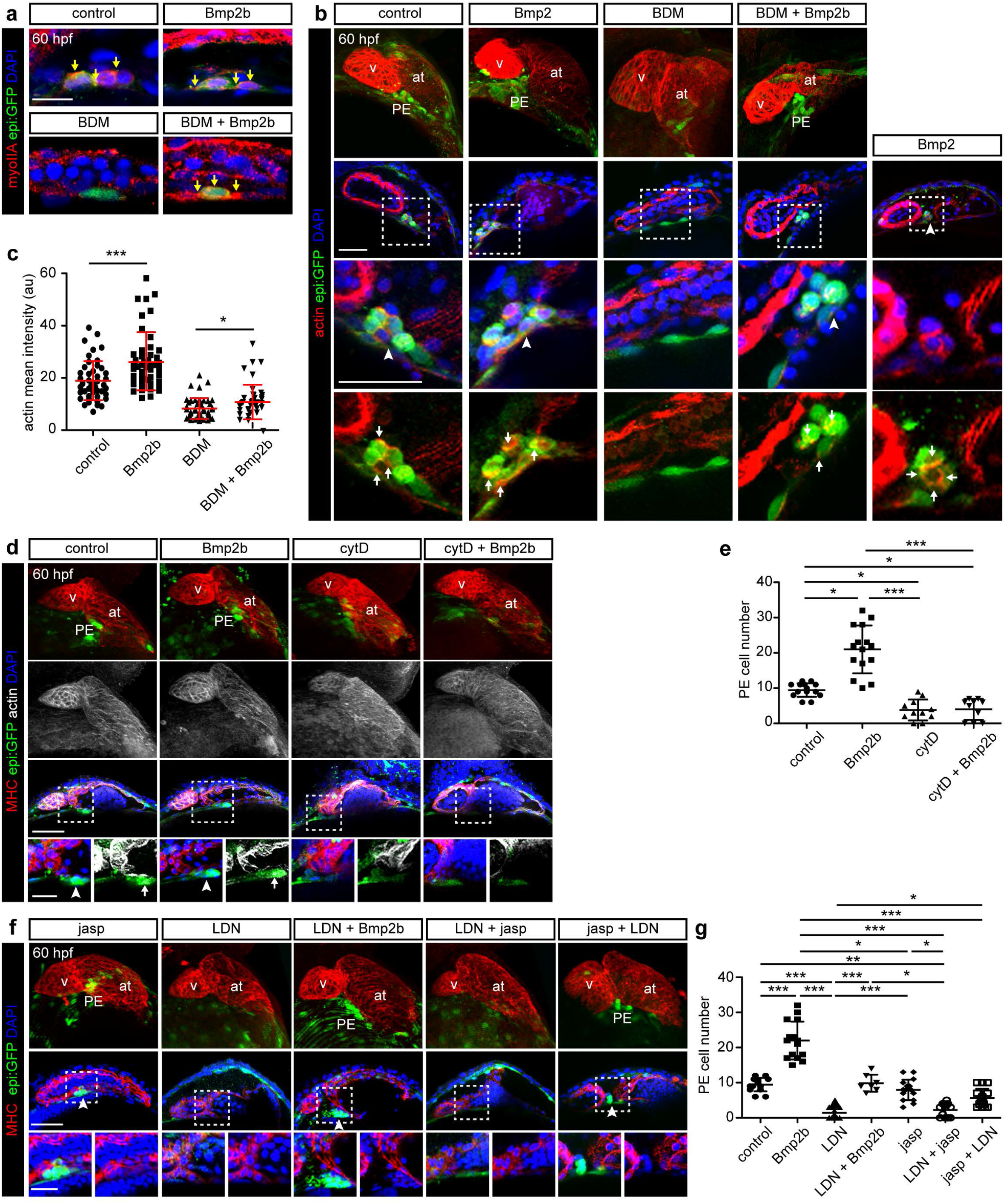
Bmp2b controls actin rearrangements necessary for PE formation. **(a)** Heart and pericardial cavity from untreated zebrafish compared with Bmp2-overexpressing and BDM-treated fish at 60 hpf. Zoomed view of the DP-PE area is shown. Pericardium and PE are in green, myosin II-A is in red and nuclei were counterstained with DAPI (blue). Yellow arrows mark myosin II-A apicocortical accumulation in PE cells. **(b)** Maximum projection of 60 hpf zebrafish hearts; optical sections of the hearts are shown in the middle panels, and zoomed images (bottom panels) are views of the PE region. Untreated fish were compared with BDM-treated and *bmp2b*-overexpressing fish. Pericardium and PE are in green and actin is in red. Nuclei are counterstained with DAPI. Arrowheads mark the PE cluster. Arrows point to actin concentration sites in the PE cluster. **(c)** Quantification of actin intensity (arbitrary units) in PE cells in **b. (d)** Top images are maximum projections of 60 hpf hearts from epi:GFP control or bmp2b-overexpressing fish treated with cytochalasin D (cytD), immunostained for GFP (green), myosin heavy chain (MHC, red) and phalloidin-Alexa488 (actin, white). Nuclei are counterstained with DAPI (blue). **(e)** Quantification of PE cell number from embryos as shown in **d. (f)** Maximum projections and optical sections of animals treated with LDN-193189 (LDN) and jasplakinolide (jasp). epi:GFP^+^ cells are in green, the MHC^+^ heart tube in red, nuclei are counterstained with DAPI (blue). **(g)** Quantification of PE cell number in **f**. at, atrium; BDM, Butanedione Monoxime; hpf, hours post fertilisation; PE, proepicardium; v, ventricle. Scale bar: 50 μm except for a (10 μm) and zoomed views in **d** and **f** (20μm). Data are mean ± s.d., one-way ANOVA followed by Kruskal-Wallis significant difference test. \**P* < 0.05, \*\**P* < 0.01, \*\*\**P* < 0.001.

Our results show that actin filaments are necessary for PE formation. In agreement with this, treatment with the actin polymerization and elongation inhibitor, cytochalasin D (cytD) at 2 μM from 48 hpf onwards, decreased the size of PE clusters from 9 ± 3 cells (in non-treated fish) to 4 ± 3 cells (n=10–14 embryos in each group) at 60 hpf. Of note, *bmp2b* overexpression was unable to rescue PE formation in the presence of cytD (Fig. 5d,e), indicating that the inhibitory effect of cytD on F-actin assembly is downstream of the effect of Bmp2b on the actin cytoskeleton in PE cells.

We next evaluated whether the effect of Bmp inhibition on PE formation could be rescued by increasing actin stabilization. Thus, the number of cells in PE clusters of animals treated with the Bmp receptor-I inhibitor LDN-193189 (LDN)^47^ was compared with those of animals treated with a combination of LDN and jasp. LDN treatment reduced the number of PE cells per cluster compared with untreated controls and reverted the increase in PE size upon *bmp2b* overexpression (Fig. 5f,g). The mean number of PE cells was 2 ± 2 cells (from a total of 24 animals) for the LDN treated group and 3 ± 2 (from a total of 9 animals) for the LDN + jasp group (Fig. 5g). In both cases, the mean number of PE cells was lower than that usually observed in untreated controls, which was 9 ± 2 cells. By contrast, when we stabilized actin filaments with jasp from 48 hpf onwards and 4 h prior to LDN administration, the number of PE cells was significantly increased to 6 ± 3 (from a total of 11 animals) compared with the LDN group. Taken together, these results show that the Bmp signaling pathway influences actomyosin polymerization and the stabilization of the F-actin network partially compensates for the negative effect of Bmp signaling inhibition on PE formation.

To gain deeper insight into the mechanisms of Bmp2b action, we analyzed how DP cell displacement is altered in BDM-treated animals in a background of *bmp2b* overexpression. We imaged epi:GFP animals from 52 to 60 hpf and tracked epi:GFP^+^ cells in the DP. In *bmp2b*-overexpressing animals, the DP constricts to the midline (Fig. 6a), as observed in controls (Fig. 1c). Upon BDM treatment, the typically observed crowding of GFP^+^ cells at the midline was not apparent (Fig. 6b). However, *bmp2b* overexpression rescued DP cell displacement towards the midline upon BDM treatment (Fig. 6c). We next quantified DP cell displacement using divergence fields. BDM treatment led to expansion, whereas control and *bmp2b* overexpression led to DP tissue constriction (Fig. 6d,e). Accordingly, cell displacement tracking revealed a movement towards the midline in control (n=4 embryos) and *bmp2b*-overexpressing embryos (n=4), whereas DP cells were predominantly displaced away from the midline in BDM-treated embryos (n=5) (Fig. 6f). Again, *bmp2b* overexpression in BDM-treated animals increased the direction of displacement and favored the movement towards midline (n=3). Altogether, the results suggest that bmp signaling acts through actomyosin cytoskeleton to allow the displacement of DP cells towards the midline, which ultimately leads to PE formation.

**Fig. 6.**
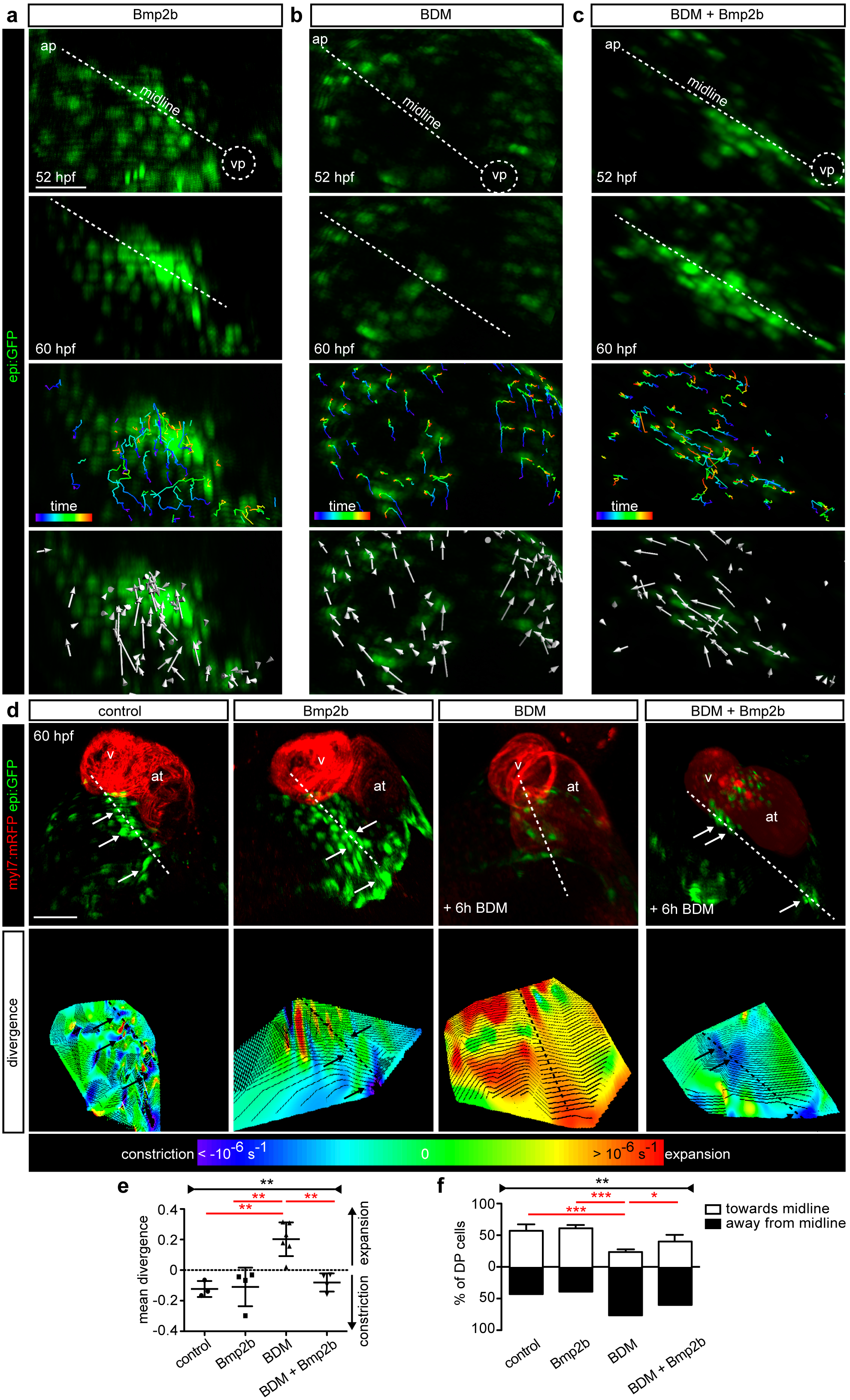
Effect of Bmp2b and myosin II inhibition on dorsal pericardial layer constriction. **(a-c)** First and last frame of an epi:GFP *in vivo* time-lapse, **(a)** bmp2b-overexpressing, **(b)** BDM-treated or **(c)** Bmp2b overexpression in BDM-treated larvae; midline is shown by a discontinuous white line. Colored tracks label first time-frame in purple and the last in red. Arrows indicate overall direction of tracked cells. **(d)** Ventral view of *epi:GFP;myl7:mRFP* hearts. Shown are maximal projections of: untreated, bmp2b-overexpressing, BDM-treated and *bmp2b-overexpression* in BDM-treated fish. Bottom images show 3D reconstruction of the divergence field of the DP; blue regions represent constriction and red-orange regions of tissue expansion. Arrows point to PE cells; dotted lines draw the midline. **(e)** Mean divergence of the tracked cells within the DP from *in vivo* time-lapses. **(f)** Percentage of cells that move towards or away from the midline. Data in **e** and **f** are mean ± s.d., one-way ANOVA followed by Kruskal-Wallis significant difference test was used and is shown with black line and asterisk the summary, ** *P* < 0.01. Unpaired two-tailed Student’s *t*-test was also used and shown with red lines and asterisks, * *P* < 0.05; ** *P* < 0.01, *** *P* < 0.001. ap, arterial pole; at, atrium; BDM, Butanedione Monoxime; hpf, hours post fertilization; PE, proepicardium; v, ventricle; vp, venous pole. Scale bar: 50 μm.

### Endocardial/endothelial Notch signaling acts upstream of Bmp2b to control PE formation

Notch and Bmp signaling pathways are connected during the development of several organs including the heart^48–52^. Moreover, loss of function of Notch signaling alters epicardium formation and Notch signaling regulates smooth muscle differentiation of epicardium-derived cells^53^. To study the link between Notch and Bmp2 signaling pathways during PE formation in the zebrafish, we used the transgenic line *UAS:NICD-myc^KCA4^*^54^ to overexpress the intracellular active domain of the Notch receptor (NICD) under a HS-inducible promoter *hsp70:Gal4^KCA3^*. When we performed HS in this line at 48 hpf, PE formation was unaltered at 60 hpf when compared with non-transgenic zebrafish (8 ± 4 cells *vs* 8 ± 3 cells, respectively) (Fig. 7a,b). However, the overexpression of *NICD* led to the maintenance of a PE as late as 80 hpf, a time point at which a PE cluster is typically no longer visible (Supplementary Fig. 4a,b) (6 ± 3 cells *vs* 0 ± 1 cells in controls; *P* < 0.0001).

**Fig. 7.**
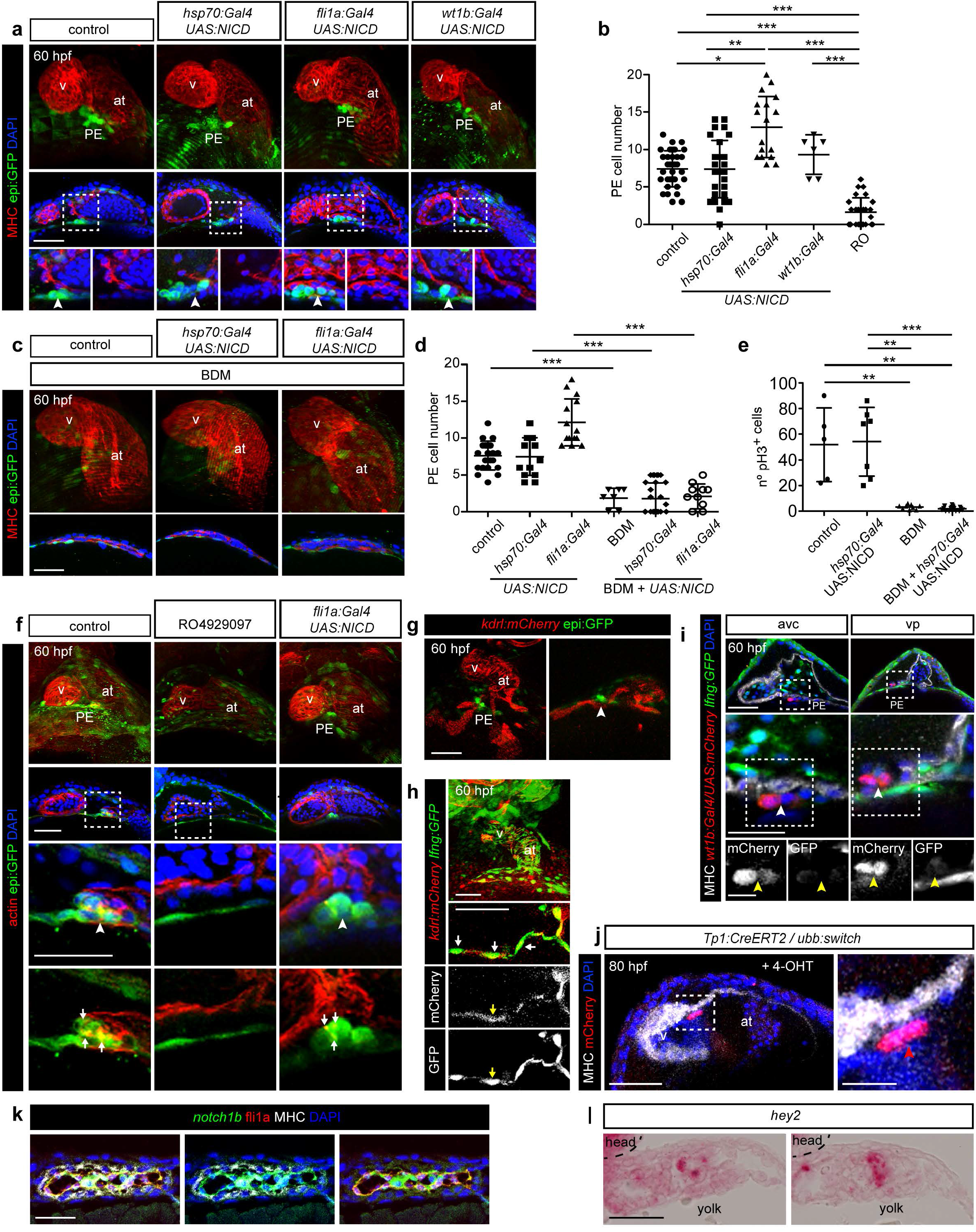
Effect of Notch signaling activation on PE formation. **(a)** Maximal projection and optical sections at 60 hpf of control embryos and embryos overexpressing NICD (*UAS:NICD*) upon heat shock induction in all cells (*hsp70:Gal4*), in endocardial and endothelial (*fli1a:Gal4*) pericardial and proepicardial cells (*wt1b:Gal4*). Immunostaining was performed for GFP (green), MHC (red) and nuclei were counterstained with DAPI (blue). Arrowheads point to the PE. **(b)** Quantification of PE cell number in **a. (c)** Maximal projection and optical sections at 60 hpf of larvae BDM treatment. **(d)** Quantification of PE cell number in **c. (e)** Quantification of pH3^+^ cells in the pericardium. **(f)** Maximum projection of 60 hpf zebrafish hearts; optical sections are shown in the middle panels, zoomed images (bottom panels) are views of the PE region. Untreated fish were compared with RO-treated animals and larvae overexpressing *NICD*. GFP labels the pericardium and PE (green), actin labeled with phalloidin is shown in red. Nuclei are counterstained with DAPI (blue). Arrows point to actin concentration sites in the PE cluster. **(g)** epi:GFP crossed with the *kdrl:mCherry* line. Maximal projection and optical sections of a representative frame from an *in vivo* time-lapse. GFP^+^ pericardium and PE are shown in green, the mCherry^+^ endocardium in red. **(h)** *lnfg:GFP* crossed with the *kdrl:mCherry* line. Maximal projection and optical section of a representative frame from an *in vivo* time-lapse. **(i)** Optical sections through the avc and vp regions of *wt1b:Gal4; UAS:mCherry; lnfg.GFP* larvae at 60 hpf. PE cells (red), *lnfg:GFP* (green), MHC^+^ heart tube (white), cell nuclei (DAPI, blue). Zoomed views including single channels are shown in the bottom panels. Arrowheads point to mCherry^+^/GFP^−^ PE cells. **(j)** *Tp1:CreERT2;ubb:switch* embryos were treated with 4-Hydroxytamoxifen to trace the fate of Notch1 responsive cells with mCherry. At 80 hpf, mCherry^+^ cells are observed in the endocardium. **(k,l)** Fluorescent *in situ* mRNA hybridization for *notch1a* (k, green, n=3/3 embryos) and RNAscope detection of *hey2* **(l)** followed by immunofluorescence for *fli1a:GFP* (red) and MHC (gray) on 60 hpf heart sections (2/2 embryos). Nuclei are counterstained with DAPI (blue). at, atrium; avc, atrio-ventricular canal; DP, dorsal pericardium; hpf; hours post fertilization; MHC, Myosin Heavy Chain; PE, proepicardium; v, ventricle; vp, venous pole. Scale bar: 50 μm except for i middle images, 20 μm and zoomed images in **k** (10 μm) and l (25 μm). Data are mean ± s.d., one-way ANOVA followed by Kruskal-Wallis significant difference test in **b** and Student’s *t*-test in **d**. *** *P* < 0.001.

To determine the cell type in which the activity of NICD is necessary, we crossed *UAS:NICD-myc^KCA4^* transgenic fish with *wt1b:Gal4* animals to drive NICD expression in DP and PE cells, or with *fli1a:gal4^ubs3Tg^* ^55^, to overexpress NICD in endothelial and endocardial cells (Fig. 7a,b). Whereas activation of the Notch pathway in *wt1b^+^* cells did not affect PE formation, its activation in *fli1a^+^* cells resulted in the significant increase in PE cell numbers, which was already apparent at 60 hpf, as well as a maintenance of the PE cluster at 80 hpf (Supplementary Fig. 4a,b) (14 ± 8 cells *vs* 0 ± 1 cells in controls; *P* < 0.0001). Therefore, Notch-over activation in endothelial/endocardial cells affects PE cluster formation. Next, we tested whether *NICD* overexpression was also able to rescue PE formation under BDM treatment, as observed for *bmp2b*. At 60 hpf, PE formation was not rescued by *NICD* overexpression (Fig. 7c,d) but at 80 hpf a PE cluster was observed in BDM-treated animals (13 ± 8 cells *vs* 0 ± 1 cells in BDM only; *P* < 0.0001;Supplementary Fig. 4c,d). *NICD* overexpression did not increase cell proliferation in the DP, as assessed by quantification of pH3^+^ cells and it also did not rescue the number of proliferating cells under BDM treatment (Fig. 7e). The effect of Notch1 inhibition for PE formation was also assessed with the Notch inhibitor, RO-4929097 (RO)^56^. RO administration reduced PE cluster size (2 ± 2 cells *vs* 8 ± 3 cells in controls), which occurred concomitant with the loss of actin cytoskeleton polarization in DP cells at the midline (Fig. 7 b,f). These findings suggest that paracrine signals from the underlying endothelium or endocardium guide PE formation. Indeed, we observed that at the site of cluster formation at the DP, the PE is situated on top of *kdrl*^+^ endothelial sinus venosus horns (Fig. 7g and Supplementary Movie 17). Moreover these endothelial cells show GFP expression in *kdrl:mCherry* transgenic fish crossed to *ET33-mi60a* animals^57^. This enhancer trap line, herein called *lfng:GFP*, expresses GFP under the control of the *lunatic fringe* (*lfng*) gene regulatory region, a modulator for Notch signaling^58,59^ (Fig. 7h). *lfng:GFP* marks endothelial cells from the sinus venosus and the endocardium (Fig. 7f,g). PE cells do not present high levels of *lnfg:GFP* expression (Fig. 7h,i). As a further proof-of-concept that Notch activity is present in endocardial cells, we lineage traced Notch responsive cells using *Tp1:CreERT2*^60^. In this line, *CreERT2* is driven by RBPK binding sites. Crossing *Tg(Tp1:CreERT2)* into *Tg(ubb:LOXP-EGFP-LOXP-mCherry)* and treating embryos with 4-hydroxytamoxifen (4-OHT) at 48 hpf allows for the permanent labeling of cells that receive Notch signaling at the time of 4-OHT administration by mCherry expression. At 80 hpf, GFP^+^ cells were detected only in the endocardial cells at the atrioventricular canal, but not in the epicardium (Fig. 7j). Fluorescent *in situ* mRNA hybridization to detect *notch1a* revealed that its coexpression colocalizes with fli1a:GFP^61^, an endocardial reporter line (Fig. 7k). Furthermore, the Notch pathway target *Hey2* is expression in the endocardium of the embryonic heart tube (Fig. 7l). Altogether our data indicate that endocardial/ endothelial Notch pathway activation is required for PE formation. The fact that the rescue of PE formation under BDM treatment by *NICD* overexpression occurred with a delay in comparison to that seen with Bmp2b activity suggests that the Notch pathway might act upstream of *bmp2b* in the control of PE formation.

To further study a possible effect of Notch on Bmp signaling during PE formation we examined the presence of PE cells and pSmad1/5+ DP cells upon endothelial/endocardial Notch overactivation in *fli1a:gal4;UAS:NICD* animals. At 60 hpf we observed that animals with ectopically-activated Notch signaling in *fli1a^+^*, but not in *wt1b^+^* cells, revealed pSmad1/5+DP and PE cells, suggesting an upregulation of Bmp signaling in pericardial cells in response to endothelial Notch activity (Fig. 8a-c). At 80 hpf, this effect is exacerbated (Supplementary Fig. 4e-g).

**Fig. 8.**
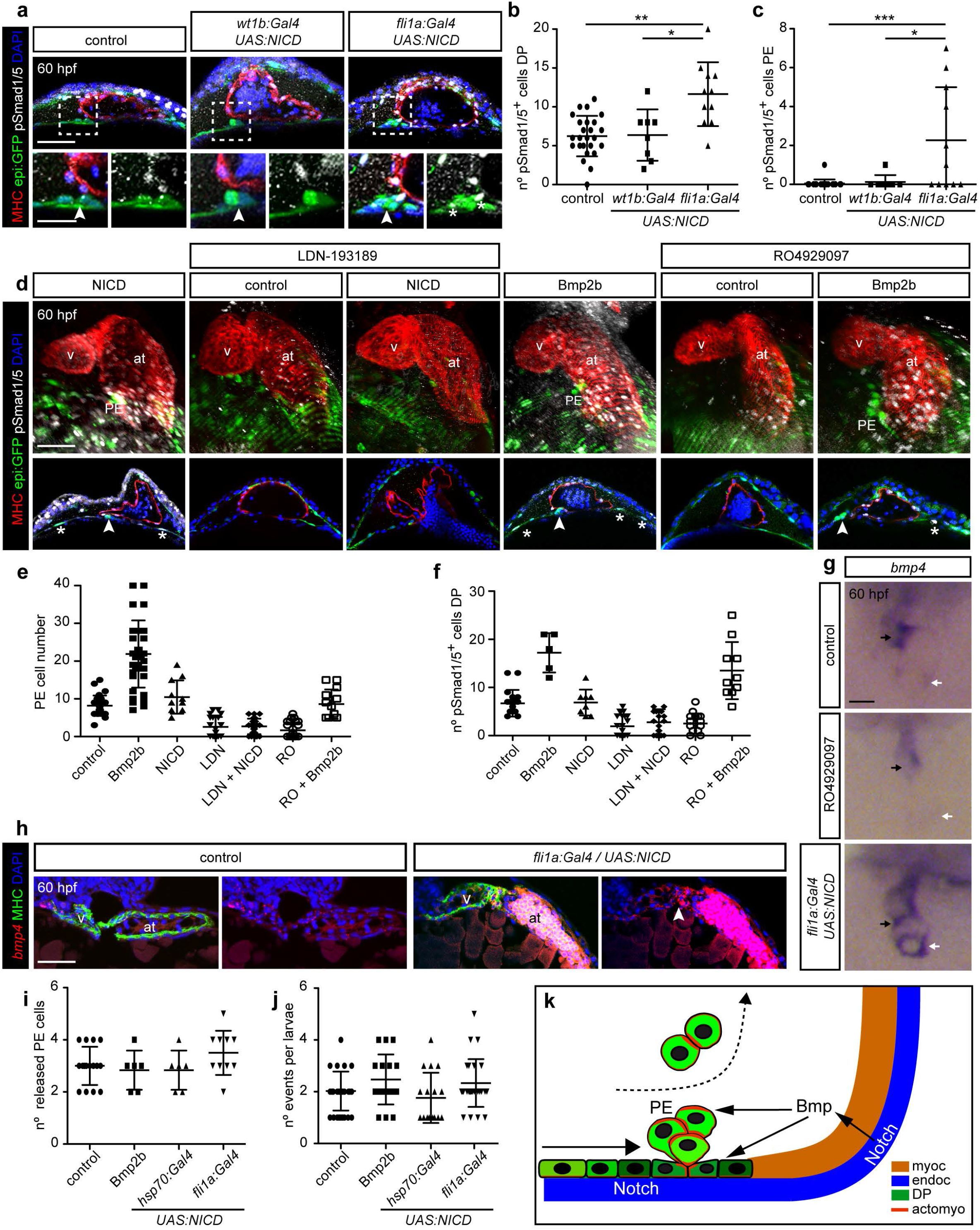
Endothelial Notch signaling acts upstream of Bmp to control PE formation. **(a)** Maximal projections and optical sections of larvae at 60 hpf overexpressing NICD in DP/PE cells (*wt1b:Gal4*) or in the endothelium (*fli1a:Gal4*). Zoomed views are shown below. epi:GFP^+^ cells are in green, MHC in red, pSmad1/5 in white and nuclei were counterstained with DAPI (blue). Arrowheads mark PE cluster and asterisks pSmad1/5^+^ PE cells. **(b)** Quantification of pSmad 1/5^+^ DP cell numbers. **(c)** Quantification of pSmad1/5^+^ PE cell numbers. **(d)** Maximal projections and optical sections of LDN193189 (LDN) or RO4929097 (RO)-treated animals at 60 hpf in non-transgenic versus *bmp2b* or *NICD*-overexpressing animals using *hsp70:Gal4*. DP and PE are in green, heart tube in red, pSmad1/5 in white. Nuclei were counterstained with DAPI. Arrowheads indicate the PE. Asterisks point to pSmad1/5^+^ DP cells. **(e)** Quantification of PE cell number as shown in **d. (f)** Quantification of DP pSmad1/5^+^ cell number in **d. (g)** Whole mount *in situ* hybridization for *bmp4* in control, RO-treated and endothelial *NICD* overexpressing animals. White arrows point to the venous pole, black arrows to the atrioventricular canal (avc) of the heart tube. **(h)** *in situ* hybridization for *bmp4* on sections from control (2/3 embryos) and endothelial *NICD* overexpressing animals through the heart. *bmp4* expression is enhanced upon *NICD* overexpression (2/3 embryos, arrowhead). **(i)** Number of PE cells shown during each event of cell release per larvae observed from 55-64 hpf. **(j)** Number of events of cell release. **(k)** Model of the interaction between Notch, BMP and the actomyosin network on PE formation. A section of a 60 hpf larvae close to the venous pole was drawn. at, atrium; DP, dorsal pericardium; hpf; hours post fertilization; HS, heat shock; MHC, Myosin Heavy Chain; PE, proepicardium; v, ventricle. Scale bar: 50 μm except for zoomed images in **a** and **h** (20 μm). Data are mean ± s.d., one-way ANOVA followed by Kruskal-Wallis significant difference test. \**P* < 0.05, \*\**P* < 0.01, \*\*\**P* < 0.001.

We also analyzed the impact of the increase in PE cells upon *NICD* and *bmp2b* overexpression on epicardium formation. Whereas epicardial cell numbers were unaffected by the overexpression of *NICD, bmp2b* overexpression increased the epicardial cell number at 80 hpf as compared with untreated animals (Supplementary Fig. 4h). Similarly, we wanted to assess whether PE formation was stable over longer time periods: at 5 days postfertilization (dpf) the PE was not present (Supplementary Fig. 4i). To further address the relationship between Notch and Bmp signaling pathways, we combined various Notch and Bmp gain- and loss-of-function scenarios. First, we added LDN from 48 hpf to heat-shocked *hsp70:Gal4^KCA3^;UAS:NICD-myc^KCA4^* embryos (Fig. 8d). At 60 hpf, PE cell formation was not rescued by *NICD* overexpression in presence of LDN (2 ± 3 cells vs 9 ± 2 cells in untreated animals) (Fig. 8e). The reduction in PE formation upon RO treatment from 48 hpf onwards was reverted by the overexpression of *bmp2b* where a cluster similar to that seen in a wildtype condition was present (2 ± 2 cells vs 8 ± 5 cells, *P* < 0.0001) (Fig. 8d,e).

Next, we evaluated how the different treatments affected pSmad1/5 levels in the DP (Fig. 8f). As expected, *bmp2b* overexpression increased the number of pSmad1/5+ cells and LDN treatment reduced this. RO administration also led to a reduction in the number of pSmad1/5+ cells and this was rescued by *bmp2b* overexpression (n=10), but HS-induced *NICD* overexpression could not rescue the number of pSmad1/5+ cells upon LDN treatment. Given that pSmad1/5 levels increased upon Notch overactivation, we analyzed whether bmp expression levels were altered in the heart tube. At 60 hpf, RO treatment reduced *bmp4* expression levels (Fig. 8g, n=22/26 for control and 12/20 for RO-treated animals). Under endothelial *NICD* overexpression, *bmp4* expression is increased in the heart tube (n=9/13). At 80 hpf, we also detected increased *bmp4* and additionally *bmp2b* expression under endocardial *NICD* overexpression, and reduced levels in RO-treated animals (Supplementary Fig. 4j, n=20 embryos for each condition). Fluorescent *in situ* hybridization against *bmp4* was performed to analyze in which cell type this gene is expressed. We found colocalization of *bmp4* expression with the myocardial marker Myosin Heavy Chain in control and endocardial *NICD* overexpression (Fig. 8h).

One possible explanation for the increase of PE clusters observed upon *bmp2b* or *NICD* overexpression is that less cells are released from the PE and hence that PE clusters change in size in the different tested conditions. Thus, we counted the number of PE cell release events during 8 h *in vivo* and no differences were observed between control, *bmp2b* or *NICD* overexpression (Fig. 8i). Similarly, the number of PE cells released per event did not differ between groups (Fig. 8j).

Overall, these experiments indicate that the Notch signaling pathway acts in the endothelium and endocardium to regulate Bmp signaling in the myocardium, which is necessary for PE formation (Fig. 8k).

## DISCUSSION

We propose a model in which Notch activity in endocardial cells leads to the expression of *bmp2b* and *bmp4* in the embryonic heart tube myocardium, which subsequently signals to PE precursor cells promoting PE cluster formation through its effect on the actomyosin cytoskeleton (Fig. 8k). These findings illustrate the importance of an intact actomyosin scaffold for generating the interactions and forces between DP cells necessary for the apical extrusion of PE cells, the source of the epicardial layer. Thus, PE formation is a highly dynamic morphogenetic event in which the combined action of mechanical events and signaling pathways converge.

At the midline of the DP, individual DP cells delaminate from the pericardial layer towards the cavity, and therefore PE cluster formation is based on apical extrusion. Apical extrusion is a cellular mechanism that has been reported to control epithelial layer homeostasis in the intestine while preserving its barrier function^62–64^. Live cell extrusion has also been reported to occur in epithelia during embryogenesis to control cell number^30^. Accordingly, an increase in cellular density leads to the elimination of supernumerary cells towards the apical side^30,65^. Extruded cells then undergo apoptosis upon loss of cell contact with their neighbors^66,67^. Neighboring wild-type cells also play an active role in this process by accumulating cytoskeleton as intermediate filaments at the interface with transformed cells^68^. Instances where extruded cells survive, as observed here during epicardium formation, have been documented for transformed cells: specifically in epithelial cancer models where apically-extruded transformed cells overexpress the oncogenes *Src* or *Ras*, which prevent cell death^69–72^.

Here we describe a physiological mechanism of apical cell extrusion that results in cell survival and is part of a natural process occurring during heart development that is required for the generation of epicardial precursor cells. To our knowledge, this is the first description of apical extrusion occurring during embryogenesis that leads to living extruded cells, but it may represent only one example of a series of morphogenetic events in which this mechanism is involved. An additional process where apical extrusion is likely to play a role is during the emergence of hematopoietic stem cells from the dorsal aorta^73,74^. The gene regulatory mechanism involved in promoting the survival of extruded PE cells, until they attach to the myocardium, remains to be elucidated. It is plausible, that rapid attachment of extruded PE cells to the myocardial surface promotes their cell survival.

Our results reveal a previously unidentified role for pericardial cell movements in PE formation. We found that these motions are dependent on the actomyosin cytoskeleton and lead to an overall constriction of the DP tissue at the midline. Interestingly, actomyosin-dependent cell rearrangements at the DP influence outflow track development in the mouse^75^.

Cell proliferation contributed partially to cell crowding at the midline and influenced PE formation. However, ectopic *bmp2b* overexpression could promote PE formation in the absence of cell proliferation, which raises the question of the extent to which cell proliferation is a limiting step for DP cell movement towards the midline. During gastrulation, cell rearrangements are not always regulated by proliferation as they can be controlled by global adherent junction-remodeling arising from myosin-II-mediated local contractile forces between cells^76^. This is reminiscent of the movement of DP cells, which we observed during PE cluster generation that also is dependent on myosin II activity.

The interplay between cell signaling and mechanics during development is a major focus of research^77^. An important question to be answered is how Bmp2 signaling and actomyosin dynamics are linked to control PE formation. Enhanced Bmp2b signaling could overcome the negative effect of BDM on PE formation by blocking myosin II activity. Our results suggest that Bmp2b counteracts the tissue compliance caused by inhibition of the actomyosin network. They could also indicate that Bmp2b stabilizes F-actin: since actin intensity levels were higher upon Bmp2 overexpression, a higher dose of BDM would be required to disrupt the actomyosin network. Another possibility is that Bmp2 rescues the F-actin tension by modulating the expression of other non-conventional myosins that are not inhibited by BDM. Indeed, Bmp signaling has been recently shown to regulate epithelial morphogenesis by controlling F-actin rearrangements, apical constriction, cell elongation and epithelial bending^78^. Along this line, interkinetic nuclear migration during morphogenesis of the retina is driven by actomyosin forces that are blocked when myosin II is inhibited, and the reduction in the velocity of migration and cytoskeleton dynamics by BDM can be rescued by BDM-insensitive myosins^79^. Furthermore Bmp2 has been shown to control the expression of the *non-muscle myosin Va* gene, thereby promoting cellular migration^80^. Bmp2 signaling has been recently suggested to regulate *Prrx1* expression in the lateral plate mesoderm, which in turn regulates the expression of *palladin*, an actin bundling protein^27^. This signaling cascade might be important for PE formation.

Our results suggest a direct effect of Bmp2b signaling on actomyosin dynamics, but it is also conceivable that Bmp2b acts on DP cells by changing their cell-cell adhesion or cell-basal membrane interactions, which would allow them to overcome the absence of a functional actomyosin network. E-cadherin and tight junction-associated proteins are involved in polarization and apical extrusion of transformed epithelial cells^81^. Integrin-paxillin signaling also plays an important role in PE formation^12^, and Bmp signaling has been previously shown to affect integrin signaling. Bmp2 can induce spreading of a myoblast cell line by activating integrins, which increases cell adhesion dynamics and leads to reorganization of the cytoskeleton^82^. Moreover, integrins *per se* are Bmp targets^83^. Thus, a possible scenario can be envisaged in which ectopic *bmp2* promotes PE formation by promoting integrin-basement membrane interactions. Studies determining how the Bmp pathway affects the actomyosin cytoskeleton and whether it relates to the redistribution in junction components such as E-cadherin or integrins during PE formation will help to reveal the process of PE cell detachment from the DP.

Bmp signaling has been described to be necessary for PE specification in the zebrafish^9^. In chicken PE explant assays, a balance between Bmp and Fgf signaling determines the specification of precardiac mesoderm into either PE or cardiomyocytes^84,85^. Furthermore, Bmp is one of the factors needed to drive differentiation of human pluripotent stem cells into an epicardial lineage^86,87^. A relationship between Notch and Bmp2 signaling during cardiovascular development has already been reported in the mouse^51,88^. *N1ICD* gain of function experiments showed that myocardial Notch1 overexpression negatively regulates Bmp2^52^. Consistently, Notch signaling abrogation leads to increased Bmp2 signaling in the PE^53^. Thus, whereas the Notch/Bmp2 axis promotes PE formation in the zebrafish, it represses it in the mouse. In mammals, the PE is formed by an outer mesothelial layer and an inner mesenchymal core^89^, whereas in the zebrafish, PE cells are derived from the pericardial mesothelium^14^. It is possible that this distinct PE architecture contributes to a different effect of Notch and Bmp2 signaling.

In conclusion, our findings illustrate the importance of an intact actomyosin scaffold for generating the interactions and forces between DP cells necessary for the apical extrusion of PE cells, the source of the epicardial layer. Collectively, our results reveal an exquisite orchestration between heart tube maturation and PE formation and may represent a paradigm for the coordinated action of signaling molecules and mechanical forces in controlling tissue morphogenesis.

## MATERIALS AND METHODS

### Zebrafish strains and husbandry

All experiments were approved by the Community of Madrid “Dirección General de Medio Ambiente” in Spain; and the “Amt für Landwirtschaft und Natur” from the Canton of Bern, Switzerland. Animals were housed and experiments performed in accordance with Spanish and Swiss bioethical regulations for the use of laboratory animals. Fish were maintained at a water temperature of 28 °C. The following fish were used: wild-type AB strain; Et(−26.5Hsa.WT1-gata2:EGFP)^cn1^ (epi:GFP)^14^; *Tg(myl7:mRFP)*^90^, *Tg*(*hsp70l:bmp2b*)^fr13^ and Tg(*hsp70l::noggin3*)^fr14^^45^, Tg(*kdrl::mCherry*)^ci5^ (from Elke Obers laboratory), Tg(*uas::myc-Notch1a-intra*)^kca3^, Tg(*hsp70l::Gal4*)^kca4^^91^; Tg(*fli1a:gal4*)^ubs3Tg^^55^, Tg(*βactin:LifeActin:RFP*)^e2212Tg^^92^, Tg(*actb2:myl12.1-mCherry*)^e1954^ ^93^, Tg(*BRE-AAVmlp:dmKO2*)^mw40^^46^, Et(*krt4:EGFP*)^sqet33mi60A^ ^57^, Tg(*Tp1:CreERT2*)^s951^ ^60^, Tg(-*3.5ubb:LOXP-EGFP-LOXP-mCherry*)^*cz1702*^ ^94^ Tg(*fli1a:eGFP*)^61^ and Tg(*UAS.mRFP*)^95^.

The Et(*-26.5Hsa.WT1-gata2:EGFP*)^cn1^ line contains a reporter construct flanked by FRT sites, in which cardiac actin drives the expression of RFP. The cassette was removed by injection of flipase into one-cell stage embryos. We named this new line Et(*-26.5Hsa.WT1-gata2:EGFP*)^cn14^.

For experiments to overexpress NICD in the endocardium/endothelium the following triple transgenic line was used: Tg(*fli1a:gal4*);(*hsp70:gal4*);(*UAS:NICD*) and the control was Tg(*hsp70:gal4*);(*UAS:NICD*), without the HS step.

### Heat shock

Heat shock was performed to the embryos at 39°C in preheated water for 1 h.

### Generation of the *TgBAC*(*wt1b:GAL4*) transgenic line

The translational start codon of *wt1b* in the BAC clone CH73-186G17 was replaced with the *galff-polyA-Kan^R^* cassette by Red/ET recombineering technology (GeneBridges) as described^96^. To generate the targeting PCR product, *wt1b*-specific primers were designed to contain 50 nucleotide homology arms around the ATG with ~20 nucleotide ends to amplify the *galff-polyA-Kan^R^* cassette To facilitate transgenesis, the BAC-derived *loxP* site was replaced with the *iTol2-Amp^R^* cassette^97^ using the same technology. The final BAC was purified with the HiPure Midiprep kit (Invitrogen) and co-injected with Tol2 mRNA into *Tg*(*UAS:GFP*) embryos^95^. The full name of this transgenic line is *TgBAC*(*wt1b:GALFF*).

Primers used to generate the *wt1b*-GALFF targeting PCR product were wt1b_HA1_Gal4-For: gacattttgaactcagatattctagtgttttgcaacccagaaaatccgtcaccATGAAGCTACTGTCTTCTATCGAAC and wt1b_HA2-KanR-Rev: gcgctcaggtctctgacatccgatcccatcgggccgcacggctctgtcagTCAGAAGAACTCGTCAAGAA (lower case indicates homology arms).

### Immunofluorescence

Embryos were fixed overnight in 4% paraformaldehyde in PBS, washed in 0.01% PBS-Tween-20 (Sigma) and permeabilized with 0.5% Triton-X100 (Sigma) in PBS for 20 min. Several washing steps were followed by blocking for 2 h with 5% goat serum, 5% BSA, 20 mM MgCl2 in PBS followed by overnight incubation with the primary antibody at 4°C. Secondary antibodies were diluted 1:500 in PBS and incubated for 3 h. Nuclei were counterstained with DAPI (Invitrogen) for 30 min. After several washes, embryos were mounted in Vectashield (Vector).

The antibodies and stains for immunofluorescence detection were as follows: anti-myosin heavy chain (MF20, DSHB) at a 1:20 dilution, anti-pH3 (Millipore) at 1:100, anti-GFP (aveslab) at 1:1000, anti-pSmad1/5 (Cell Signaling Technology) at 1:100, Phalloidin-488 (Thermo Fisher) at 1:100, anti-myosin IIA (Sigma) at 1:100 and anti-mRFP (Abcam) at 1:250. Secondary antibodies were the following: anti-mouse-Cy3 (The Jackson Laboratory), anti-mouse IgG2b 568 (Invitrogen), anti-chicken 488 (Life Technologies), anti-rabbit 647 (Thermo Scientific), all diluted at 1:500.

Embryos were imaged with a Zeiss 780 confocal microscope fitted with a 20× objective 1.0 NA with a dipping lens. Z-stacks were taken every 3–5 μm. Maximal projections of images were 3D reconstructed in whole-mount views using IMARIS software (Bitplane Scientific Software). The pericardial ventral tissue was digitally removed to provide a clearer view of the heart. Optical sections of 1–3 z-slices were also reconstructed.

### Quantification of DP and PE cells

PE cells have been described to emerge from two main regions of the DP: the avcPE appears close to the atrio-ventricular canal, and the vpPE around the venous pole. We counted each cell in each z plane using DAPI and GFP expression using the line epi:GFP. We took care not to count any cell twice. Cells with a round morphology at the vp or avc region were counted as PE cells, Cells with a flat morphology in the DP were counted as DP cells. See Supplementary Fig. 5 and Supplementary Movie 18 for further explanation.

### Actin mean intensity measurement

Images from 3 embryos of each condition at 60 hpf were acquired at the same conditions with a Zeiss 780 confocal microscope. Z-stacks were taken every 5 μm. Acquired images were in monochrome 8-bits. Z-slides where the PE was present were opened in ImageJ (NIH) and squared areas of 3.32 × 3.32 pixels were drawn in the channel of the PE cluster. The mean intensity was calculated for actin (phalloidin staining) in this PE region. The intensity brightness sum was normalized to the number of pixels in the selected area to obtain the actin mean intensity brightness in the PE. Brightness was in arbitrary units from 0–255 (where 0 is a pixel with no staining pixel and 255 a saturated pixel). Mean intensity values were represented as spots in a histogram.

### Pharmacological treatments

Embryos were manually dechorionated and incubated with compounds from 48 hpf onwards (unless otherwise stated). The following compounds were used: aphidilcolin (150 μM), hydroxyurea (20 μM), nocodazole (0.01 mg/ml), BDM (10–20 mM), LDN-193189 (20mM), cytocalasin D (2 μM) (all from Sigma), BLEB (25–50 μM; Abcam), jasplakinolide (0.15 μM; Thermofisher), RO-4929097 (10 μM; Selleckchem), 4-OHT (5 μM; Sigma).

### In situ hybridization

ISH on whole mount embryos was performed as described^98^ using riboprobes against full coding sequence of *bmp4* or *bmp2b* cDNAs as well as *notch1b*^99^. Embryos at 60 hpf or 80 hpf were fixed in 4% PFA overnight, dehydrated in methanol series and stored at −20°C until its use. On day 1, embryos was bleached in 1,5% of H_2_O_2_ in methanol, rehydrated, washed in TBS with 0.1% Tween20 (TBST), digested with proteinase K 10 μg ml^−1^ for 17 min, rinsed in TBST, blocked the endogenous alkaline phosphatase with 0,1M triethanolamine pH 8 with 25 μl/ml of acetic anhydride for 20 min, washed in TBST, re-fixed in 4% PFA for 20 min. After washing again in TBST they were pre-hybridized at 68°C for at least 1 h. The antisense riboprobe was added at 0.5 μg ml^−1^. After overnight hybridization, two washed with 50%Formamide/5xSSC plus 2% Tween20, four washed with 2xSSC plus 0,2% Tween20, all at 68°C. Then, embryos were transferred to RT, washed in TBST and incubated with 10% heat inactivated goat serum, 1,2% of blocking reagent (Roche, 11096176001) in maleic acid buffer (MABT). Then, embryos were incubated overnight with 1:4000 dilution of anti digoxigenin-AP antibody (Roche, 11093274910) in blocking solution. After overnight, embryos were washed in MABT and developed in BM-Purple until signal was detected.

ISH on paraffin sections were done following commercial RNAscope protocol (Advanced Cell Diagnostics)^100^.

For fluorescent *in situ* hybridization combined with immunostaining in paraffin sections were deparaffinized, post-fixed 20 minutes with PFA 4%, washed with PBS, treated with proteinase K 10 μm ml^−1^ for 10 minutes at 37 °C, washed with PBS, post-fixed with PFA 4% for 5 minutes, washed with PBS, treated with HCl 0.07 N for 15 minutes, washed with PBS, treated with 0.25% acetic anhydride in triethanolamine 0.1 M pH 8 for 10 minutes, washed with PBS, washed with RNase free water and then hybridized with the probe in pre-hybridization buffer over night at 65 °C. The following day sections where washed twice with post-hybridization buffer 1 (50% Formamide, 5xSSC, 1% SDS) for 30 minutes at 65 °C and twice more with post-hybridization buffer 2 (50% Formamide, 2xSSC, 1% SDS). Then, sections were washed with MABT buffer at room temperature, and incubated at least 2 hours in blocking solution at room temperature. They were next incubated overnight with anti-digoxigenin-POD antibody (1:500) in blocking solution. The third day, they were washed with MABT for several hours, after that sections were incubated with 1:200 tyramides (Perkin-Elmer, NEL701A001KT), washed with PBST. Following with the immunofluorescence incubated the sections with Myosin Heavy Chain antibody (DSHB hybridoma bank, MF20 1:20) overnight at 4°C. The fourth day washed the slices and incubated with secondary antibody and counterstained with DAPI.

### In vivo imaging

Embryos were transferred to fish water containing 0.2 mg/ml tricaine (Sigma) and 0.0033% 1-phenyl-2-thiourea (Sigma), and immobilized in 0.7% agarose (NuSieve GT Agarose, Lonza) in a 35-mm Petridish with a glass cover (MatTek Corporation). Zebrafish hearts were scanned bidirectionally at 30 frames per second (fps) with an SP5 confocal microscope (Leica) using a 20× glycerol immersion objective with 0.7 NA. Videos were acquired every 5 μm, with a line average of 6 and a pinhole of 1.9 AU. Around 65 z-stack videos were acquired per heart every 10-15 min. GFP, red and brightfield channels were acquired simultaneously.

High resolution *in vivo* imaging was performed with the Zeiss LSM880 airy scan fastmode, using a 40x/1.1 water immersion objective lens. Sampling was performed with 1x Nyquist Coefficient parameters. Airy scan processing was performed in ZEN 2.3.

Realigned 4D data sets were displayed and analyzed using Imaris (Bitplane AG) or ImageJ.

### Drift correction

Small movements of the plate and the embryo in the agarose were manually suppressed using a drift correction, selecting a particle, usually a noise voxel, following the movement to be corrected in two consecutive frames. Moreover, a posterior drift correction was applied to suppress the intrinsic movement of the developing embryo. Microscope and growth drift was corrected in Imaris. The venous pole was identified as a stable reference structure and tracked in each frame. Subsequently, the “Correct drift” tool was applied to the resulting track.

### Cell tracking

Cell tracking was performed using a built-in Imaris tool, allowing for an automatic creation of 4D trajectories of dorsal pericardial cells. A region of interest was manually selected and after applying a background subtraction algorithm – Gaussian filter – an autoregressive algorithm is applied to perform cell tracking. Such autoregressive algorithm works under the assumption that particles move from frame to frame in a quasi-predictive fashion, interpolating the data from previous frames. This approach is pertinent in cells embedded in a tissue and hence was chosen. 4D trajectories were further filtered out in terms of duration and overall displacement to remove false-positive trajectories that would add noise to further calculations.

### Divergence of velocity field calculator tool

The Algorithm to obtain the divergence of the dorsal pericardium velocity field was determined as follows: Divergence of the velocity field associated with the movements of the dorsal pericardium was calculated as an indicator of the expansion of the tissue. The divergence is a mathematical operator that, when applied to a velocity field, indicates how much a set of particles expand or constrict in space. In this particular case, the set of particles is given by the dorsal pericardial cells, hence giving an estimate of the expansion/constriction of the tissue and characterizing it kinematically. It is mathematically described in Equation 1.

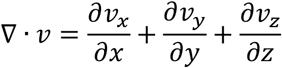

To do so, a customized Matlab code was created. Taking as an input the 4D trajectories calculated with Imaris a 2D grid was created, which was used to interpolate the z-position of the dorsal pericardial cells, hence resembling the geometry of the dorsal pericardium. The spacing of the grid satisfies the Nyquist criteria, such that Δx_grid_ < 2*size of the cell nucleus. This interpolation was calculated using a Delaunay triangulation method. In a similar procedure, each velocity component was interpolated across the mesh using a spline method. The interpolated velocity field and z-position were used to calculate the divergence of the velocity field.

Since the divergence calculated takes into account 3 dimensions, but the geometry of the tissue is a laminar 2D surface embedded in a 3D space, 2D divergence were taken in the XY, YZ, XZ planes and algebraically operated to obtain the 3D divergence as described in Equation 2. 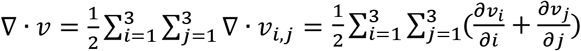 (Equation 2). After calculation of the divergence field each node of the z-interpolated surface is assigned a divergence value, with positive values indicating an expansion and negative values a constriction.

Additionally, the code provides the mean divergence of the surface for each time frame, according to Equation 3. 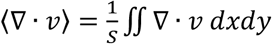 (Equation 3).

### Angle calculation and directionality to midline

To calculate the angle of cell movement to the midline, epi:GFP embryos were *in vivo* imaged from 52 hpf. The midline is a defined line that runs from the venous pole to the outflow tract of the heart tube. Dorsal pericardium epi:GFP^+^ cells were tracked using Imaris software during the time-lapse, and the movement vector was calculated regarding the first and the last position of the cell. The angles that form the cell tracks vectors with the midline were calculated in ImageJ. The directionality of the cell tracks in relation to the midline was calculated with regards to the relative distance of the first and last cell track point to the midline vector. Distance of first track point to midline > distance of last track point = cell movement towards midline; distance of first track point to midline < distance of last track point = cell movement against midline.

### Statistical analysis

Student’s t test for comparisons between two groups or one-way ANOVA for comparisons between more than two groups was used when normal distribution could be assumed. When the normality assumption could not be verified with a reliable method, the Kruskal-Wallis test was used. Model assumptions of normality and homogeneity were checked with conventional residual plots. The specific test used in each comparison is indicated in the figure legend. Calculations were made with Microsoft Excel and GraphPad. P-values are indicated either in the figure legends, the main text or summarized in Supplementary Table 1.

## ACKNOWLEDGMENTS

We are grateful to the Animal facility and Microscopy Units from CNIC, and the Anatomy, Histology and Neuroscience department from the UAM, MIC-Bern and J. Langa and R. Baal for fish husbandry of the University of Bern. We thank Mathias Hammerschmidt, Salim Seyfried, Carl-Philipp Heisenberg and Nikolay Ninov for sharing transgenic lines. N.M. was funded by the Spanish Ministry of Economy and Competitiveness through grant BFU2014–56970–P (Plan Estatal de Investigación Científica y Técnica y de Innovatión 2013-2016. Programa Estatal de I+D+i Orientada a los Retos de la Sociedad Retos Investigación: Proyectos I+D+i 2016, del Ministerio de Economía competitividad e Industria), and co-funding by Fondo Europeo de Desarrollo Regional (FEDER). N.M. is also supported by the Swiss National Science Foundation grant ANR-SNF 310030E-164245, the ERC starting grant 337703 and European Industrial Doctorate Program EID 722427. L.A-D was funded through the postdoctoral fellowship Ayudas Postdoctorales 2013. JL de la Pompa was supported by grants SAF2016-78370-R, CB16/11/00399 (CIBER CV) and RD16/0011/0021 (TERCEL) from the Spanish Ministry of Economy, Industry and Competitiveness (MEIC). The CNIC is supported by the Ministry of Economy, Industry and Competitiveness (MEIC) and the Pro CNIC Foundation, and is a Severo Ochoa Center of Excellence (MEIC award SEV-2015-0505).

